# Interpenetrating networks of fibrillar and amorphous collagen promote cell spreading and hydrogel stability

**DOI:** 10.1101/2024.09.11.612534

**Authors:** Lucia G. Brunel, Chris M. Long, Fotis Christakopoulos, Betty Cai, Patrik K. Johansson, Diya Singhal, Annika Enejder, David Myung, Sarah C. Heilshorn

## Abstract

Hydrogels composed of collagen, the most abundant protein in the human body, are widely used as scaffolds for tissue engineering due to their ability to support cellular activity. However, collagen hydrogels with encapsulated cells often experience bulk contraction due to cell-generated forces, and conventional strategies to mitigate this undesired deformation often compromise either the fibrillar microstructure or cytocompatibility of the collagen. To support the spreading of encapsulated cells while preserving the structural integrity of the gels, we present an interpenetrating network (IPN) of two distinct collagen networks with different crosslinking mechanisms and microstructures. First, a physically self-assembled collagen network preserves the fibrillar microstructure and enables the spreading of encapsulated human corneal mesenchymal stromal cells. Second, an amorphous collagen network covalently crosslinked with bioorthogonal chemistry fills the voids between fibrils and stabilizes the gel against cell-induced contraction. This collagen IPN balances the biofunctionality of natural collagen with the stability of covalently crosslinked, engineered polymers. Taken together, these data represent a new avenue for maintaining both the fiber-induced spreading of cells and the structural integrity of collagen hydrogels by leveraging an IPN of fibrillar and amorphous collagen networks.

**Statement of significance:** Collagen hydrogels are widely used as scaffolds for tissue engineering due to their support of cellular activity. However, collagen hydrogels often undergo undesired changes in size and shape due to cell-generated forces, and conventional strategies to mitigate this deformation typically compromise either the fibrillar microstructure or cytocompatibility of the collagen. In this study, we introduce an innovative interpenetrating network (IPN) that combines physically self-assembled, fibrillar collagen—ideal for promoting cell adhesion and spreading—with covalently crosslinked, amorphous collagen—ideal for enhancing bulk hydrogel stability. Our IPN design maintains the native fibrillar structure of collagen while significantly improving resistance against cell-induced contraction, providing a promising solution to enhance the performance and reliability of collagen hydrogels for tissue engineering applications.

**Graphical abstract:** 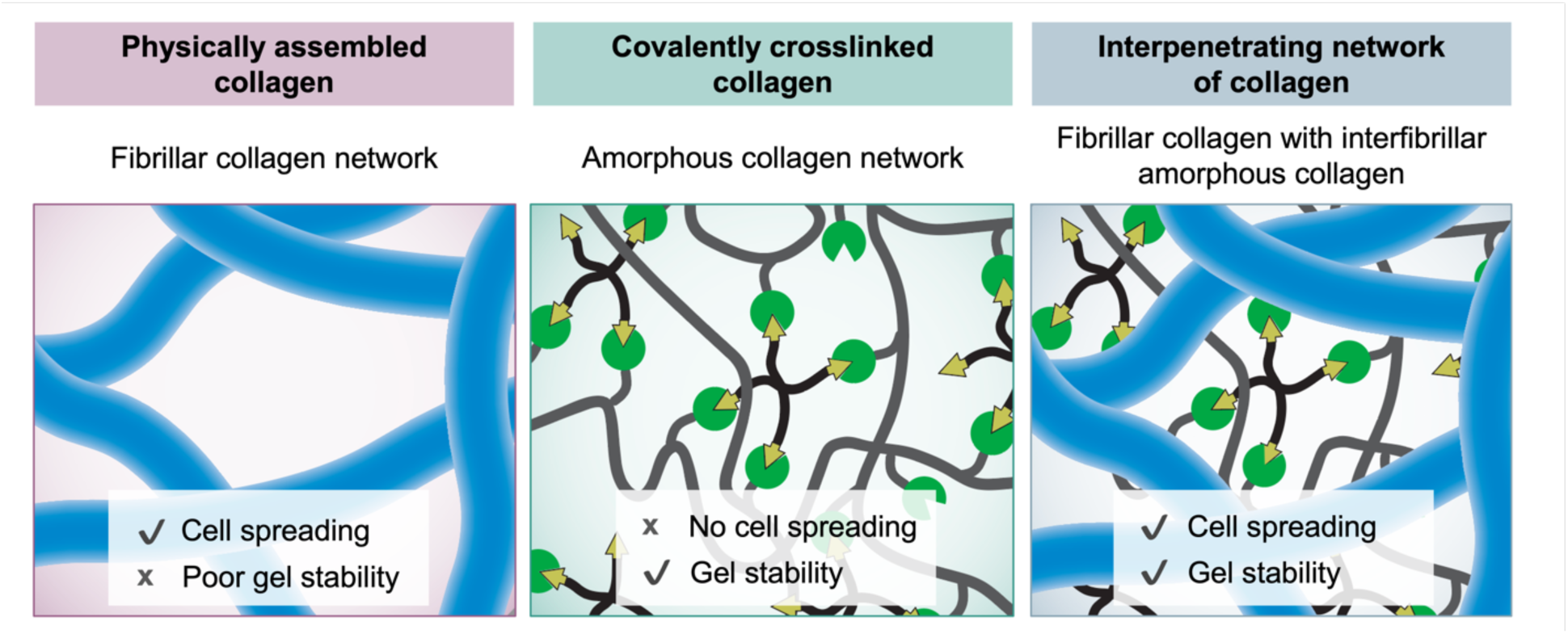

## 1. Introduction

Collagen hydrogels are widely used as three-dimensional (3D) scaffolds for tissue engineering and regenerative medicine applications due to their favorable material and cell-interactive properties. As the most abundant protein in the human extracellular matrix (ECM), collagen comprises approximately 25% of the body’s total protein content by dry weight [1]. Its inherent biocompatibility and minimal risk of immune rejection, particularly in the form of atelocollagen, make it a clinically viable material for medical implantation [2,3]. Additionally, the fibrillar structure of collagen type I and its numerous ligand-binding sites promote cell adhesion and interaction, thereby directing essential cellular behaviors including spreading, migration, and differentiation [4–6]. The ability of collagen hydrogels to closely mimic the native ECM provides a significant advantage over synthetic hydrogels, supporting more natural cellular functions and superior tissue regeneration outcomes.

However, a significant drawback of collagen hydrogels is their tendency to contract over time in response to forces from contractile, encapsulated cells that exert significant tensions on the surrounding collagen fibers as they actively remodel the matrix [7–9]. This contraction presents several challenges for tissue engineering applications. For example, it can lead to instability of the engineered gels, causing substantial shrinkage and deformation of the gel shape and size. These structural changes can disrupt the gel’s alignment with patient-specific geometries, reduce porosity, and alter cell density, which impacts cell viability, differentiation, and overall tissue function [10,11]. Additionally, gel contraction can hinder the efficacy and engraftment potential of engineered tissue by causing detachment from the intended implantation site [12]. This phenomenon has been observed across a variety of cell types, including fibroblasts and mesenchymal stromal cells, with the severity of contraction increasing with the time in culture [11,13,14].

Several strategies have previously been employed to strengthen the bulk properties of collagen gels and reduce gel deformation, each with its own limitations [15,16]. For example, increasing collagen concentration enhances the mechanical properties of the gel but also alters its fibrillar structure, resulting in more, but thinner fibers that can significantly alter cell behavior [17,18]. Further, high-density collagen limits the ability of cells to proliferate, differentiate, and diffuse waste products [19]. The crosslinking of collagen with biological covalent crosslinkers like genipin has been used; however, these crosslinkers also modify the collagen microstructure and reduce the gel’s pore size [20,21]. Chemical crosslinkers such as glutaraldehyde, 1-ethyl-3-(3-dimethyl aminopropyl) carbodiimide hydrochloride (EDC), and N-hydroxysuccinimide (NHS) are effective in forming covalent bonds to strengthen the hydrogels, but their cytotoxicity and off-target effects render them less suitable for encapsulating live cells within the hydrogels during crosslinking [16,22,23].

Strain-promoted azide-alkyne cycloaddition (SPAAC) click chemistry offers a covalent, bioorthogonal crosslinking strategy that exhibits no cross-reactivity with cells and avoids the cytotoxic catalysts, side-reactions, and harmful byproducts often associated with other crosslinking methods [24,25]. By chemically modifying the collagen to contain azide functional groups, the collagen can be crosslinked with a molecule containing strained alkynes, such as dibenzocyclooctyne (DBCO) [26]. SPAAC chemistry is not present in biological systems and thus non-interactive with cells, allowing cells to be encapsulated directly into the hydrogel precursor during crosslinking. However, unlike conventional, physically self-assembled (PHYS) collagen, SPAAC collagen does not self-assemble into a fibrillar network, a phenomenon commonly observed for chemically modified collagen [11,26–28]. Our previous studies have shown that while the covalent crosslinks in SPAAC collagen hydrogels enhance resistance against corneal cell-induced contraction, the amorphous nature of SPAAC collagen inhibits the ability of encapsulated cells to spread as they do in fibrillar microenvironments like conventional PHYS collagen [11] or the native corneal stroma [29]. As a result, new hydrogel designs are necessary to produce collagen scaffolds that provide both resistance against contraction as well as biophysical cues for cell spreading.

We hypothesized that an interpenetrating network (IPN) of collagen that incorporates both covalent crosslinking *via* SPAAC chemistry (resulting in an amorphous collagen subnetwork) and physical self-assembly (resulting in a fibrillar collagen subnetwork) would enhance hydrogel stability against cell-induced contraction while still enabling cells to spread inside the gel. IPNs are systems composed of two or more functionally distinct polymer networks interwoven at a molecular level, yet not covalently bonded to each other. These networks allow for tuning hydrogel properties to provide biomimetic microenvironments with greater control over factors like viscoelasticity. Previous studies have explored IPNs involving collagen blended with other materials such as alginate or hyaluronic acid to achieve the desired material properties [30–34]. In this work, we have developed an IPN composed of collagen with two different structural forms: fibrillar and amorphous. By forming the IPN sequentially—first by allowing physical self-assembly (PHYS collagen) and then introducing the covalent crosslinking (SPAAC collagen)—we demonstrated that the resultant fibrillar network closely resembles that of unmodified, physically self-assembled collagen. Importantly, this collagen IPN successfully facilitates cell spreading, similar to PHYS collagen alone, while maintaining structural gel integrity against cell-induced contraction, similar to SPAAC collagen alone. These properties are crucial to enable cell encapsulation for *in vitro* culture or *in vivo* implantation while preserving long-term stability and function.

## 2. Materials and Methods

### 2.1. Preparation of hydrogels

#### Preparation of PHYS collagen

Bovine type I atelocollagen solution (10 mg/mL, Advanced BioMatrix) was neutralized on ice immediately before use following instructions from the manufacturer. Briefly, 1.0 M sodium hydroxide (NaOH, Sigma), ultrapure deionized water (Millipore), and 10X phosphate buffered saline (PBS, Millipore) were added to the collagen to reach a concentration of 8 mg/mL collagen with a pH of 7.5. The solution was then diluted with cold 1X PBS to 4 mg/mL collagen. To form PHYS collagen hydrogels, the neutralized collagen solution was transferred to 8-well chambered coverslips (Ibidi, 200 μL per well) and incubated at 37°C for 1 h for gelation.

#### Preparation of SPAAC collagen

Bovine type I atelocollagen solution (10 mg/mL, Advanced BioMatrix) was modified with azide functional groups using N-hydroxysuccinimide (NHS) ester chemistry to react with primary amines on collagen. First, the acidic collagen solution was neutralized on ice following instructions from the manufacturer. Azido-PEG_4_-NHS ester (BroadPharm) was dissolved in dimethyl sulfoxide (DMSO, Fisher) at a concentration of 100 mg/mL and added to the neutralized collagen solution at 2 molar equivalents relative to primary amines on the collagen. The solution was mixed well, rotated for 2 h at 4 °C, and then dialyzed overnight in a Slide-A-Lyzer dialysis kit (3.5-kDa MWCO, ThermoScientific) against 1X PBS at 4 °C. The degree of functionalization was determined using a 2,4,6-Trinitrobenzene Sulfonic Acid (TNBSA) assay (Thermo Scientific) to quantify free amino groups, following the instructions from the manufacturer. Collagen-azide was stored at 4 °C and used within one week of the bioconjugation reaction. To form SPAAC hydrogels, collagen-azide and polyethylene glycol-dibenzocyclooctyne (PEG-DBCO, 4-arm, 10 kDa, Creative PEGworks) solutions were mixed together to reach concentrations of 2 mg/mL and 4 mg/mL, respectively. The mixture was transferred to 8-well chambered coverslips (Ibidi, 200 μL per well) and incubated at 37°C for 1 h for gelation.

#### Preparation of the collagen IPN

Neutralized, unmodified collagen solution and collagen-azide solution were mixed to final concentrations of 4 mg/mL and 2 mg/mL, respectively. For simultaneous PHYS and SPAAC network formation, PEG-DBCO crosslinker dissolved in 1X PBS was added at an effective concentration of 4 mg/mL, and the solution was then transferred to 8-well chambered coverslips (Ibidi, 200 μL per well) and incubated at 37 °C for 1 h. For sequential PHYS and SPAAC network formation, the collagen mixture without crosslinker was transferred to 8-well chambered coverslips and incubated at 37 °C for 1 h. Then, 100 μL of PEG-DBCO crosslinker dissolved in 1X PBS was added to the top of each well for an effective concentration of 4 mg/mL PEG-DBCO.

### 2.2. Mechanical characterization

Mechanical testing was performed using an ARG2 stress-controlled rheometer (TA Instruments). All measurements were confirmed to be within the linear viscoelastic regime of the materials, and representative curves were shown based on measurements of at least N = 3 independent samples.

Tests comparing PHYS and SPAAC collagen alone were conducted with a cone and plate geometry, using a 20 mm diameter and 1° angle cone, and the materials were allowed to gel *in situ* on the rheometer. All solutions were initially kept on ice, and a solvent trap or mineral oil was used to prevent evaporation during the measurement. The solutions for PHYS and SPAAC collagen were mixed at their final concentrations and immediately pipetted onto the rheometer stage. The gelation kinetics were evaluated through a temperature sweep from 4 to 37 °C (heating rate of 3 °C/min, 1 rad/s angular frequency, 1% strain) followed by a time sweep. Once the final plateau modulus was reached, a stress relaxation test (10% strain, 37 °C) was conducted.

Tests comparing PHYS, SPAAC, and IPN collagen (Figure 4) were performed with an 8-mm parallel plate geometry, and the gels were formed before loading onto the rheometer. Specifically, 4 mg/mL unmodified collagen (for PHYS collagen), 2 mg/mL collagen-azide and 4 mg/mL PEG-DBCO (for SPAAC collagen), or 4 mg/mL unmodified collagen and 2 mg/mL collagen-azide (for IPN collagen) were pipetted into silicone molds (8 mm diameter, 40 µL material) and allowed to gel for 1 h at 37 °C. For the IPN collagen, PEG-DBCO dissolved in PBS was added atop the gels for a final concentration of 4 mg/mL PEG-DBCO in the gel and incubated again for 1 h at 37 °C. Gels were removed from their molds with a spatula and placed onto the rheometer stage for frequency sweeps (0.1-100 rad/s, 1% strain, 37 °C) and stress relaxation (10% strain, 37 °C) measurements.

### 2.3. Cell culture and encapsulation

Corneal mesenchymal stromal cells (MSCs) were isolated from human donor corneas (Lions Eye Institute for Transplant and Research) according to established protocols [35]. Cells were expanded in growth medium consisting of 500 mL MEM-Alpha (Corning), 50 mL fetal bovine serum (Gibco), 5 mL GlutaMax (Gibco), 5 mL non-essential amino acids (Gibco), and 5 mL antibiotic-antimycotic (Gibco). Growth medium was changed every other day, and corneal MSCs were passaged upon reaching 80 % confluency. The corneal MSCs were used for experiments between passages 7-9. For cell encapsulations, the corneal MSCs were trypsinized, counted, pelleted, and re-suspended at a density of 5 x 10^5^ cells/mL in 4 mg/mL unmodified collagen (for PHYS collagen), 2 mg/mL collagen-azide and 4 mg/mL PEG-DBCO (for SPAAC collagen), or 4 mg/mL unmodified collagen and 2 mg/mL collagen-azide (for IPN collagen). The solutions were then pipetted into silicone molds (4 mm diameter, 10 µL material) or 18-well glass bottom chamber slides (Ibidi) (5.7 mm x 6.1 mm, 75 uL material) and allowed to gel for 1 h at 37 °C. After gelation, growth medium (for PHYS and SPAAC collagen) or growth medium with dissolved PEG-DBCO for a final concentration of 4 mg/mL PEG-DBCO in the gel (for IPN collagen) was added atop the gels. The medium was changed every other day during the duration of the culture period (5 days).

### 2.4. Microscopy of collagen fibrils

The fibrillar collagen networks were visualized using second harmonic generation (SHG) microscopy with an inverted microscope (Nikon, Ti2-E equipped with a C2 confocal scanning head and a Nikon CFI Apochromat TIRF 100XC oil immersion objective). The C2 scanner was augmented with a slidable mirror (Optique Peter) allowing switching between confocal fluorescence (with laser diode wavelengths 405, 488, 561, and 647 nm) and nonlinear imaging modalities. In nonlinear imaging mode, the SHG signal was generated by probing the samples with a picosecond-pulsed laser from a system (APE America Inc., picoEmerald S with 2 ps pulse length, 80 MHz repetition rate, and 10 cm^-1^ bandwidth) consisting of a 1031 nm mode-locked ytterbium fiber laser and an optical parametric oscillator (OPO) tunable between 700-960 nm. The OPO wavelength was set to 797 nm and the backscattered SHG signal (at a wavelength of 398.5 nm) was separated using a set of optical filters (BrightLine 400/12 bandpass, BrightLine 390/18 bandpass, Thorlabs FESH0500 shortpass) and detected pixel-by-pixel with a photomultiplier tube (Hamamatsu, R6357). The excitation power at the sample was 35 mW.

Gels were removed from their molds and imaged in 2-well chambered coverslips (Ibidi, 1.5H cover glass thickness) with 500 μL of PBS to prevent gel desiccation. For each sample, at least 9 image z-stacks were acquired in different areas of the gel. Each z-stack comprised of 21 slices spaced 1 μm apart, for an effective stack depth of 20 μm. To avoid boundary effects, acquired z-stacks were centered at least 30 μm deep into the gels. All images were collected at a resolution of 1024 × 1024 pixels (75.3 × 75.3 nm/pixel) with 10.8 μs/pixel dwell time.

### 2.5. Image analysis of collagen fibrils

The SHG image z-stacks were analyzed using a combination of the FIJI distribution of ImageJ [36] and a modified version of a MATLAB script developed by Rossen et al [37]. The z-stacks were processed using an east shadows filter and 3D Gaussian blur filtering (σ = 1). The collagen fibrils were thresholded from the background signal using Otsu’s method [38]. The resultant binary mask was then multiplied by the original processed image to effectively isolate the fibrils from the background. To fibril thickness was determined using the “Local Thickness” plugin in FIJI. The contour length, persistence length, network mesh size, and number of fibrils were determined using a MATLAB fiber-finding algorithm.

The fiber-finding algorithm used to calculate collagen fibril contour length, persistence length, network mesh size, and the number of fibrils per image was adapted from Rossen et al. [37] and implemented in MATLAB (version R2023b). This algorithm functions by (1) tracing fibrils stepwise by iteratively searching local subvolumes from a starting point until a stopping factor is reached that separates signal from void, (2) blotting out the trace to prevent repeat counting, and (3) restarting at the next starting point until all fibrils have been exhausted. A three-dimensional Gaussian point spread function (PSF) with a voxel density of 24 × 24 × 100 nm^3^ was generated and rescaled to match the input data voxel resolution (75.3 × 75.3 × 909 nm^3^). The residuals matrix of the input data and the PSF were used for tracing fibrils due to improved clarity over the raw input data. Using iterative parameterization, a length prioritization array of [30 15 10], cone angle of 30°, and stop factor of 0.5 were found to most accurately identify fibrils for these samples and were therefore used for all analyses.

### 2.6. Microscopy of SPAAC collagen network

To visualize the SPAAC collagen network, all microscopy of fluorescently labeled SPAAC collagen was performed using fluorescence imaging with an inverted microscope (Nikon, Ti2-E equipped with a C2 confocal scanning head and a Nikon CFI Apochromat TIRF 100XC oil immersion objective).

To fluorescently label SPAAC collagen in acellular gels, the SPAAC and IPN collagen gels were first prepared as described in Section 2.1. After gelation, the gels were incubated in 30 μM Alexa Fluor 488-DBCO (Click Chemistry Tools) in PBS containing 1% BSA for 1 h at 37 °C. Unreacted dye was removed by washing the gels 4 times with PBS. Gels were kept covered with aluminum foil to protect them from photobleaching before fluorescence imaging.

To fluorescently label SPAAC collagen in corneal MSC-laden gels, collagen-azide was conjugated with Alexa Fluor 647-NHS ester (ThermoFisher Scientific). 100 µg Alexa Fluor 647-NHS ester was dissolved in 10 µL DMSO and added to 500 µL collagen-azide. The solution was covered with aluminum foil to prevent photobleaching, mixed well, and rotated for 24 h at 4 °C. The solution was dialyzed in a Slide-A-Lyzer dialysis kit (7-kDa MWCO, ThermoScientific) for 3 days against 1X PBS at 4 °C to remove unreacted dye. After dialysis, this fluorescently labeled collagen-azide was mixed with non-fluorescent collagen-azide at a 1:5 ratio. This dilution of fluorescently labeled collagen-azide was then used for cell encapsulation within SPAAC and IPN collagen gels as described in Section 2.3.

### 2.7. Microscopy and image analysis of cells within gels

Cell viability and cell aspect ratio were determined from microscopy performed with a STELLARIS 5 confocal microscope (Leica) combining fluorescence and confocal reflectance imaging modalities with a 10X air objective or 40X oil immersion objective. To assess the viability of the corneal MSCs on Days 0, 1, 3, and 5 after encapsulation, Live/Dead staining was conducted using calcein AM and ethidium homodimer-1 (Life Technologies), following the manufacturer’s instructions. The numbers of live and dead cells were determined using FIJI, and cell viability was calculated as the number of live cells divided by the total number of cells. To quantify the cells’ aspect ratio (*i.e.* ratio of major to minor axis lengths) over time as an indication of cell spreading, confocal z-stack images of live cells (calcein AM-positive) were analyzed using CellProfiler software. The aspect ratio of each cell was determined by applying the two-class Otsu thresholding, hole removal, and watershed algorithms, followed by the removal of objects with surface areas less than 50 µm^2^. Three z-stack images (12 slices with 10 µm spacing) taken in different regions of the sample were analyzed for each sample for both cell viability and cell aspect ratio. All images were acquired at a resolution of 1024 × 1024 pixels (180.4 × 180.4 nm/pixel).

Hydrogel contraction imaging was performed with an epifluorescent microscope (Leica Microsystems, THUNDER Imager 3D Cell Culture) with a 2.5X air objective in bright field mode. To track the contraction of the cell-laden hydrogels over 5 days, images of the gels within their original molds were taken at each time point, and their areas were measured with FIJI.

Images of the cell nuclei and F-actin were taken on an inverted microscope (Nikon, Ti2-E equipped with a C2 confocal scanning head and a Nikon CFI Apochromat TIRF) with a 100XC oil immersion objective. On Day 5 after cell encapsulation in the gels, samples were prepared for fluorescence microscopy by fixation with 4% paraformaldehyde for 30 min, followed by 3 PBS washes for 15 min each. Cell membranes were permeabilized with 0.25% Triton X-100 in PBS (PBST) for 1 h. Nuclei and F-actin were stained by incubation with 4′,6-diamidino-2-phenylindole (DAPI, 1 μg/mL, Thermo Fisher Scientific) and phalloidin–tetramethylrhodamine B isothiocyanate (phalloidin-TRITC, 0.2 μg/ml, Sigma-Aldrich) in PBST for 1 hour. Staining was followed by three 20-min washes in PBST.

### 2.8. Statistical analysis

Statistical analyses were performed using GraphPad Prism (Version 9.5). The Shapiro-Wilk test was used to test for normality of data. For comparison of the amine quantity in unmodified vs. azide-modified collagen, the statistical analysis was performed with an unpaired t test. For comparisons of collagen fibrillar network microstructures, a one-way analysis of variance (ANOVA) with Tukey’s multiple comparisons test was used to determine statistical significance of differences in contour length, persistence length, network mesh size, fibril density, and fibril width between collagen material conditions. For comparisons of cell aspect ratio, the Kruskal-Wallis nonparametric test was used with Dunn’s multiple comparisons correction. In all cases, N ≥ 3 independent gels for each condition, and p < 0.05 was considered as statistically significant. Data are presented as mean ± standard deviation unless specified otherwise.

## 3. Results

### 3.1. Design of an interpenetrating network (IPN) of amorphous and fibrillar collagen

While collagen is a highly desirable material for 3D cell culture scaffolds, the low-stiffness fibrillar network that results from physical self-assembly makes collagen susceptible to bulk deformation and shrinkage in response to cell-generated forces. While SPAAC crosslinking of collagen mitigates this issue by increasing hydrogel stiffness and stability, amorphous SPAAC collagen on its own is not conducive to the spreading of encapsulated cells in the manner of unmodified PHYS collagen (**Figure 1**). To explore this phenomenon, we prepared PHYS collagen gels by neutralizing bovine type I atelocollagen at 4 mg/mL, encapsulating human corneal cells within the collagen solution, and allowing the cell-laden collagen to incubate at 37 °C for 1 h for complete gelation and self-assembly of the fibrillar network (**Figure 1A**, **Figure S1A**). We observed that the fibrillar network in PHYS collagen facilitates cell spreading. However, PHYS collagen is also susceptible to shrinkage during culture as cell-imposed forces remodel the collagen fibrils, causing them to rearrange and densify around the cells over time (**Figure S2**). This remodeling process propagated to significant changes in the bulk shape and size of the hydrogels (Figure 1A).

**Figure 1.**
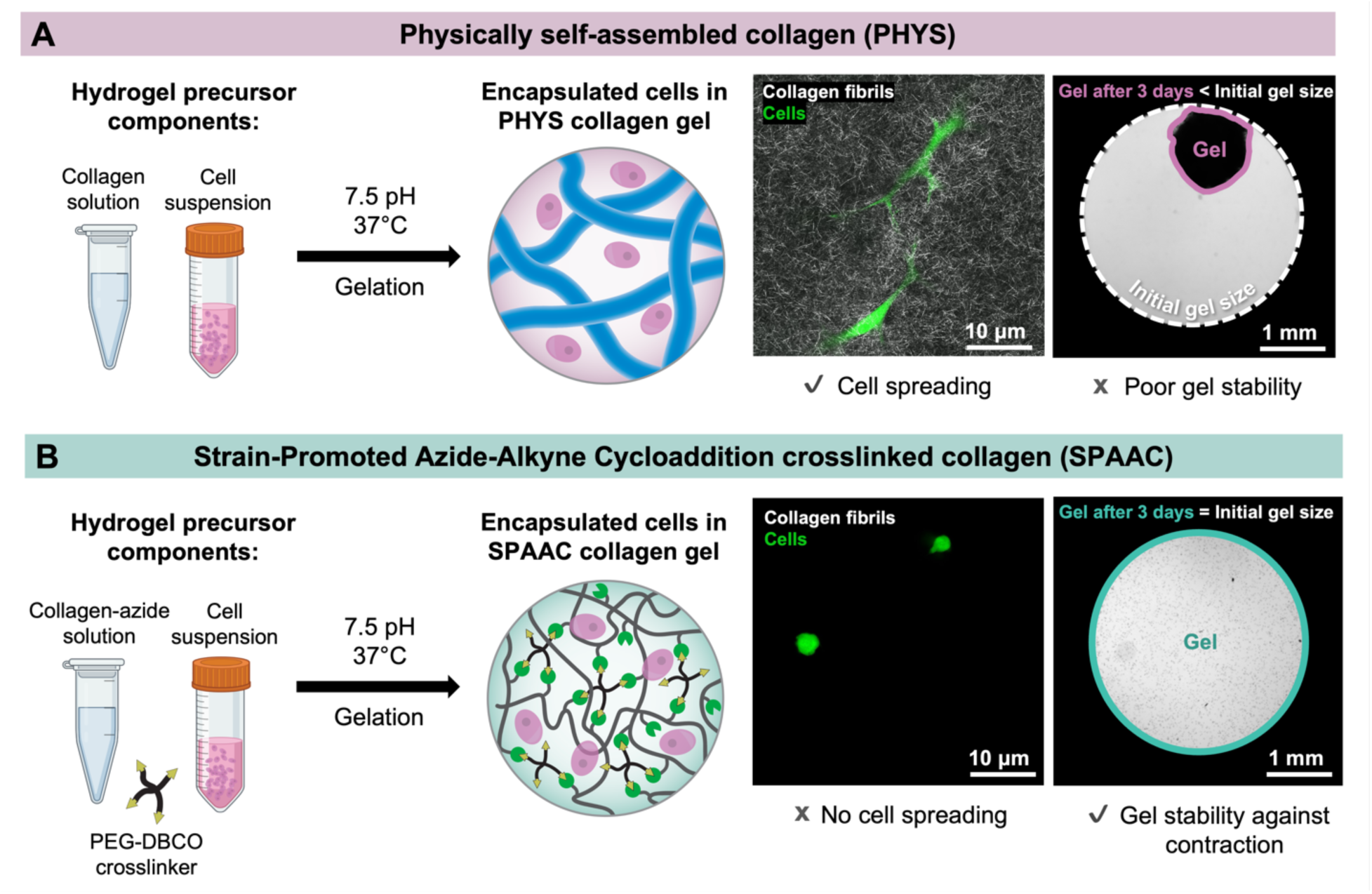
Collagen hydrogels crosslinked through physical self-assembly (PHYS) or bioorthogonal covalent chemistry (strain-promoted azide-alkyne cycloaddition, SPAAC) each offer both advantages and challenges for their use as matrices for encapsulated cells. **(A)** PHYS collagen matrices are fibrillar and facilitate the spreading of encapsulated cells, but are susceptible to contraction in response to the forces exerted by encapsulated cells. **(B)** SPAAC collagen matrices do not contract in response to the forces exerted by encapsulated cells, but do not contain fibrils or allow cell spreading.

For the SPAAC-crosslinked collagen hydrogels, we chemically modified bovine type I atelocollagen to produce collagen-azide by converting approximately 50% of the amine groups to azide groups that can participate in SPAAC crosslinking (**Figure S3**). This collagen-azide was mixed with a 4-arm PEG-DBCO crosslinker and cells (**Figure 1B**), leading to spontaneous chemical crosslinking and gelation on a timescale similar to the PHYS collagen (crossover point when G’ = G’’ in < 10 min, **Figure S1B**). A formulation of 2 mg/mL collagen-azide and 4 mg/mL PEG-DBCO was chosen to achieve a shear storage modulus (G’, representing stiffness) similar to that of the PHYS collagen gels (G’ ∼ 100 Pa, Figure S1). The covalently crosslinked SPAAC network imparted significant structural stability to the hydrogels, preventing cell-induced shrinking and compaction. However, the cells were unable to spread effectively within the SPAAC collagen hydrogels, retaining a rounded morphology (Figure 1B).

To address the challenge of balancing cell spreading and structural stability in collagen hydrogels, we developed a novel IPN that integrates two distinct structural components made from collagen: (1) a fibrillar network to support cell spreading and (2) an amorphous, covalently crosslinked network to stabilize the gel by inhibiting the long-range fibril rearrangement that causes bulk-scale contraction (Figure 2). This IPN was formulated by combining the PHYS and SPAAC collagen networks (final concentrations of 4 mg/mL unmodified collagen, 2 mg/mL collagen-azide, 4 mg/mL PEG-DBCO), creating a unique hybrid hydrogel.

**Figure 2.**
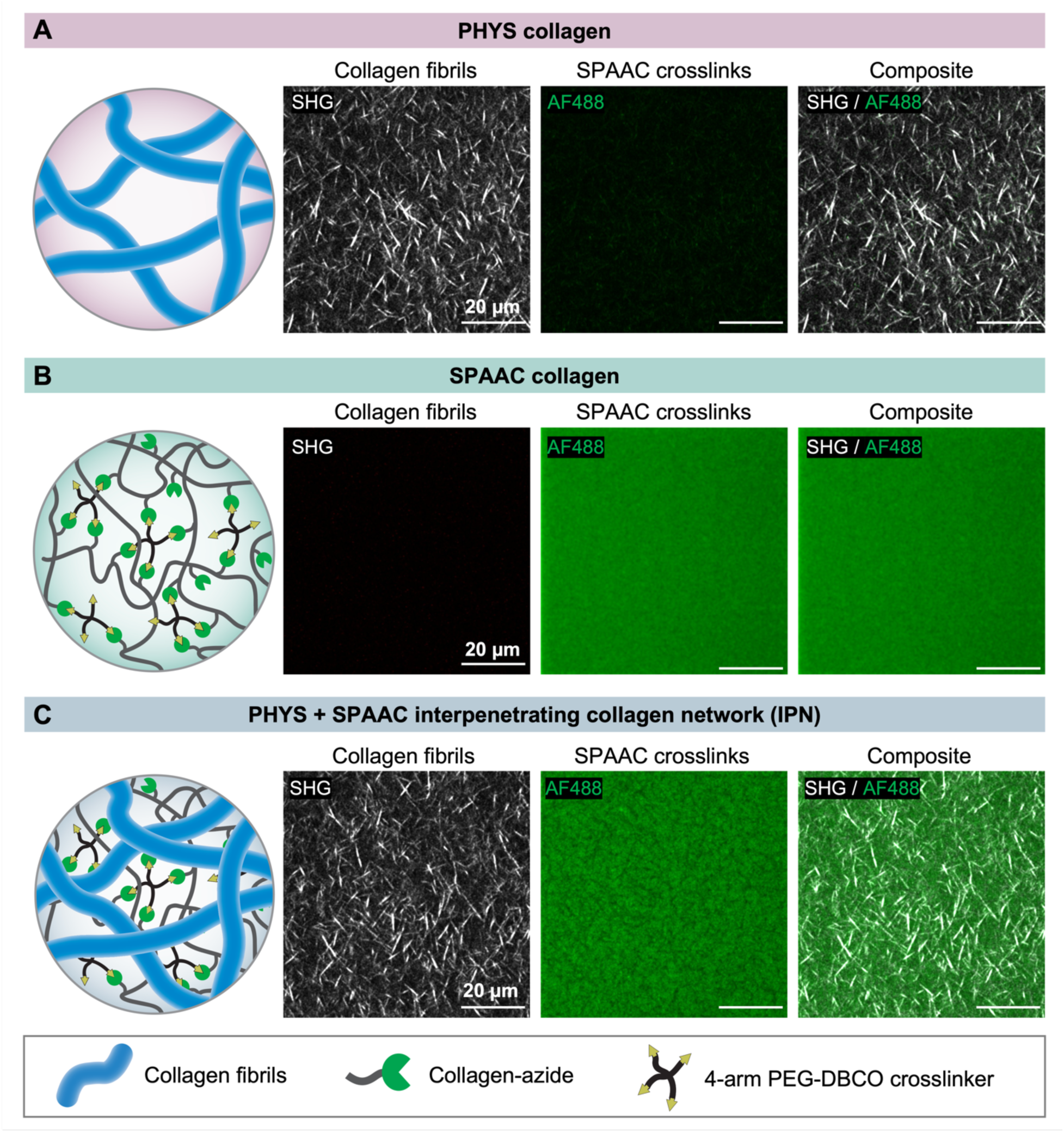
Structural characteristics of the PHYS, SPAAC, and IPN collagen materials. Collagen fibrils were visualized using second harmonic generation (SHG) imaging, and the SPAAC networks were visualized using fluorescence microscopy of fluorophore-tagged collagen-azide. **(A)** PHYS collagen possesses a fibrillar network but no covalent SPAAC network. **(B)** SPAAC collagen possesses a covalent SPAAC network but no fibrillar network. **(C)** The collagen IPN contains both a fibrillar network and a covalent SPAAC network. These two signals do not appear to overlap, which suggests the fibrillar PHYS collagen and amorphous SPAAC collagen form two distinct networks.

In its unmodified, conventional form, PHYS collagen forms a fully fibrillar network, characterized by a mesh-like arrangement of collagen struts with open voids (Figure 2A) that traditionally facilitates cell spreading. We characterized the fibrillar network in these hydrogels using second harmonic generation (SHG) imaging. This technique takes advantage of the non-centrosymmetric structure and long-range order of collagen fibrils, allowing us to visualize the fibrillar architecture through a nonlinear optical interaction that produces a strong, characteristic signal. Through this SHG signal, we confirmed the presence of a well-formed fibrillar structure in PHYS collagen. In contrast, SPAAC collagen forms an amorphous, homogenous covalent network, as visualized by tagging pendant azide groups with the fluorophore Alexa Fluor 488 (AF488) after SPAAC crosslinking (Figure 2B). The chemical modification of the amines of lysine groups, which are crucial for collagen self-assembly [39], inhibits the formation of fibrils for collagen-azide, resulting in the observed amorphous structure.

By integrating PHYS collagen and SPAAC collagen together, we created a collagen IPN featuring two distinct types of collagen crosslinking and microstructures. This IPN consists of a fibrillar PHYS collagen network embedded within an amorphous SPAAC collagen network, where the SPAAC collagen completely fills the interfibrillar spaces between the PHYS collagen (Figure 2C). When visualizing the PHYS collagen network using SHG imaging of fibrils and the SPAAC collagen network using fluorescence microscopy of AF488-tagged collagen-azide, we observed that the fluorescence signal for the SPAAC collagen was seemingly excluded from the SHG signal for the fibrillar PHYS collagen. We hypothesized that the biorthogonality of SPAAC enables the two collagen networks to be crosslinked independently in the IPN, unlike other chemistries that may result in partial grafting of the two networks. In contrast to SPAAC collagen alone (Figure 2B), the SPAAC collagen in the IPN is no longer uniformly distributed due to the presence of intervening fibrils of PHYS collagen. The spatial separation of the PHYS and SPAAC collagen networks suggests that they remain independent of each other within the IPN to form a composite structure with both fibrillar and amorphous networks coexisting.

### 3.2. Microstructure and mechanical properties of the collagen IPN

The IPN structure of this collagen hydrogel allows for modulation of the constituent subnetworks with control over their respective contributions to the material properties. However, because the PHYS and SPAAC collagen gels are formed by different mechanisms, they can interfere with each other during network formation. Both networks gelate under similar conditions, including a neutral pH of 7.5 and a physiological temperature of 37 °C. Given that the kinetic rates and resultant gelation times of the two networks within the IPN are similar (Figure S1), competition between these gelation processes can occur such that the final microstructure of the material is impacted (Figure 3). Specifically, the formation of the covalently crosslinked SPAAC collagen network can impede the self-assembly of the fibrillar PHYS collagen network. This competitive interaction results in the formation of fewer, shorter, and thinner fibrils when both networks are allowed to form simultaneously, compared to the fibrillar network observed in PHYS collagen alone (Figure 3A).

**Figure 3.**
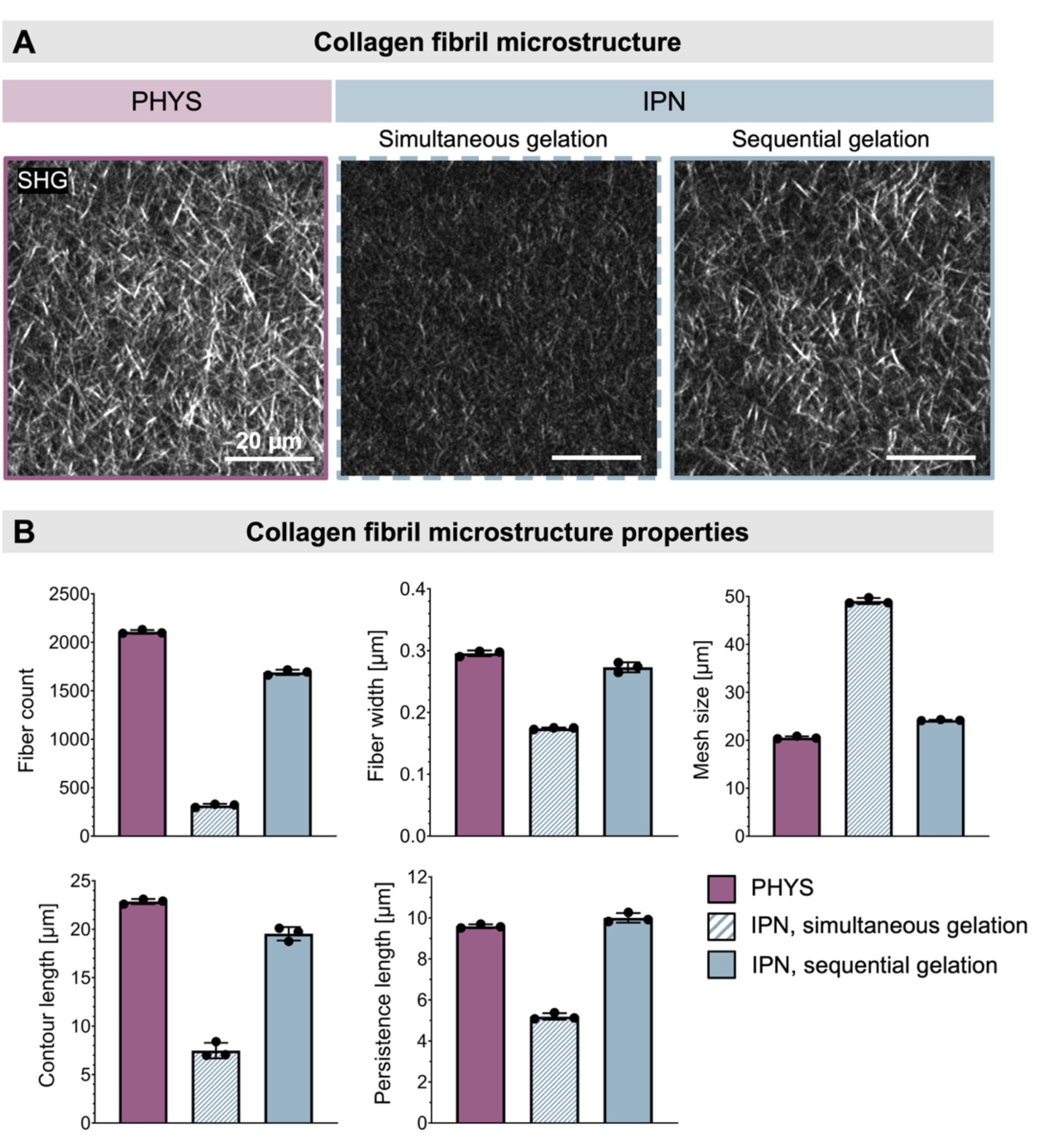
Microstructural characteristics of the PHYS and IPN collagen hydrogels. **(A)** Representative SHG imaging of the PHYS collagen, the IPN collagen with simultaneous gelation of the PHYS and SPAAC networks, and the IPN collagen with sequential gelation of the PHYS and SPAAC networks. **(B)** The IPN collagen with simultaneous formation of the PHYS and SPAAC collagen networks has fewer, thinner, and shorter collagen fibrils than PHYS collagen alone. The IPN collagen with sequential formation of the PHYS network first, followed by adding the PEG-DBCO to form the SPAAC network second yields fibrils that more closely resemble those in PHYS collagen alone. N = 3 independent gels per material condition. Data plotted as mean ± SD.

While PHYS collagen spontaneously begins to self-assemble into a fibrous network under physiological conditions, the timing of chemical crosslinking in SPAAC collagen can be controlled by adjusting when the PEG-DBCO crosslinker is added to the solution. We hypothesized that by forming the two constituent networks of the IPN sequentially, we could prevent the interference observed in collagen fibril formation when the networks are formed simultaneously. This approach is grounded in the principles of IPN synthesis, which can occur either simultaneously or sequentially. Sequential IPNs are created either by swelling a monomer or linear polymer and crosslinker into an already-polymerized single-polymer network or by selectively crosslinking one network before the other [40–42]. The method of sequential formation offers superior tunability and control over the hydrogel’s final structural characteristics and mechanical properties [43].

In our approach to creating a collagen IPN, the PHYS collagen component is first allowed to physically self-assemble at neutral pH and 37 °C for 1 h in the presence of collagen-azide. After 1 h, the PEG-DBCO crosslinker is added to initiate SPAAC crosslinking of the collagen-azide. This method ensures that the fibrillar parameters—*i.e.*, contour length, persistence length, fibril width, fiber density, and network mesh size—are maintained at levels more similar to those found in PHYS collagen alone (Figure 3B, **Table S1**). Although the presence of collagen-azide molecules, which do not participate in self-assembly into fibrils, has a minor effect on these properties relative to PHYS collagen alone, the overall fibrillar microstructure is largely maintained. Since this strategy of sequential network formation proved effective to preserve the fibrillar network, all subsequent work with the collagen IPN was conducted using this method to ensure consistent and optimal material properties.

To quantify the microstructural properties of the fibrillar hydrogels—the PHYS collagen and the collagen IPN—we used SHG imaging. The resultant SHG images provided high-resolution visualization of the fibrillar architecture, which we quantitatively analyzed to assess the effects of our sequential crosslinking method on fibril formation compared to the simultaneous crosslinking method and PHYS collagen alone. For this analysis, we employed a combination of the FIJI distribution of ImageJ and a modified version of a fiber-finding MATLAB algorithm originally developed by Rossen *et al.* [37].

This fiber finding algorithm functions by tracing lines of high-intensity pixels within a defined cone, following each line until it reaches the endpoint of a fibril. Once a fibril is fully traced, it is digitally blotted out, and the process repeats iteratively until all fibrils in the image have been characterized (**Figure S4**). To optimize this process for our particular material system and images, we fine-tuned several key parameters within the algorithm, including the cone angle, the length-prioritization array, and the pixel intensity stop factor. These modifications aid in achieving accurate fibril characterization, as they avoid pathological behavior such as identified fibrils performing sharp turns at intersections or being erroneously broken up into smaller segments. Following iterative optimization, we visually confirmed that the algorithm most accurately reflected the fibrillar structure present in our collagen hydrogels (**Figure S5**).

As expected, the differences in the microstructural properties of the SPAAC, PHYS, and IPN collagen networks also result in differences in their bulk rheological properties (Figure 4**, Figure S6**). The stiffness and viscoelasticity of hydrogels are important for tissue engineering due to their effects on cell-matrix interactions and the phenotypes of encapsulated cells [44–46]. In our system, the formulations of the constituent subnetworks of the IPN (*i.e.*, PHYS and SPAAC collagen) were selected to have a similar stiffness, as described previously: G’ ∼ 100 Pa as measured by *in situ* rheology during the gelation process (Figure S1). To allow for mechanical characterization of IPN collagen gels formed through sequential crosslinking of the PHYS and SPAAC collagen networks, we next conducted shear rheometry on gels that were pre-formed in molds, such that the PEG-DBCO crosslinker could be diffused into the IPN collagen gels. Similar to our previous results, the PHYS and SPAAC collagen gels alone had a similar stiffness to each other, although the magnitude was somewhat lower than with *in situ* gelation, likely due to the gel handling required to transfer the pre-formed gel onto the rheometer stage (Figure 4A). The stiffness of the IPN collagen, which effectively combines the PHYS and SPAAC collagen subnetworks, was greater than PHYS and SPAAC collagen alone.

**Figure 4.**
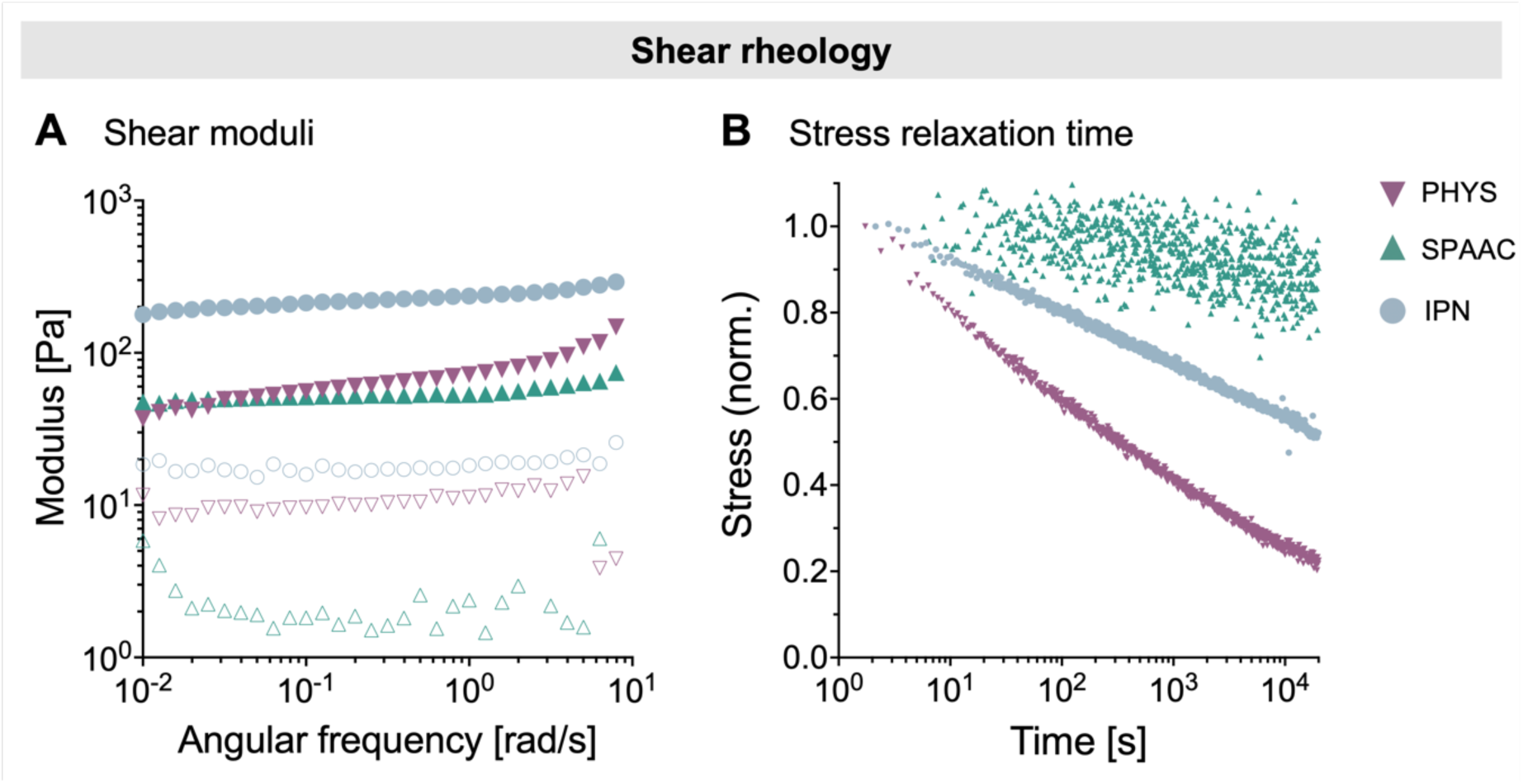
Representative mechanical characterization curves for PHYS, SPAAC, and IPN collagen gels. **(A)** The constituent components of the IPN collagen, which are PHYS and SPAAC collagen, exhibit similar shear moduli to each other. When combined into the IPN network, the shear modulus increases. Filled symbols represent the storage modulus (G’), and open symbols represent the loss modulus (G”). **(B)** PHYS collagen is rapidly stress-relaxing, while SPAAC collagen does not readily stress relax. The IPN collagen relaxes slower than PHYS collagen alone but faster than SPAAC collagen alone.

Gels exhibit stress relaxation when their polymer networks reorganize in response to an applied strain, thereby dissipating the internal stress over time. Conventional collagen gels exhibit characteristic stress-relaxing behavior due to the rearrangement of fibrils allowing for mechanical yielding and matrix flow [47–50]. Consistent with this characterization, we observed that PHYS collagen exhibits rapid stress relaxation (Figure 4B). The stress relaxation half-time (τ_1/2_, time for the initial peak stress to relax to half its original value) was approximately 300 s for PHYS collagen. This rapid stress-relaxing behavior was similar across multiple concentrations of PHYS collagen (**Figure S6A**), highlighting the role of the fibrillar, physically self-assembled microstructure in allowing dissipation of stress in the collagen network. In contrast, SPAAC collagen behaves like a mostly elastic gel due to the permanent, covalent crosslinks that do not allow for dynamic reorganization of the polymers (τ_1/2_ ∼ ∞) (Figure 4B); this behavior was also observed across multiple concentrations of SPAAC collagen gels (**Figure S6B**). When the two subnetworks of PHYS and SPAAC collagen are combined together, the collagen IPN exhibits an intermediate stress relaxation behavior (τ_1/2_ ∼ 20,000 s), which is slower stress relaxing than PHYS collagen alone but faster stress relaxing than SPAAC collagen alone.

### 3.3. Stability of collagen IPN against cell-induced contraction

Native tissues in the human body are active and dynamic biomaterials, experiencing constant interactions between the living cells and the non-living polymeric matrix in which the cells are embedded [51–53]. In particular, cells can exert forces and remodel the matrix as they undergo key cellular processes such as spreading, proliferation, and migration [33,54,55].

Cell-laden collagen hydrogels are relevant for many tissue applications including the skin, muscle, and connective tissues [56,57]. The human corneal stroma, for example, is composed of 71% collagen by dry weight, predominantly type I collagen [58]. This collagen matrix is crucial for the cornea’s biomechanical properties, and it provides a suitable extracellular microenvironment for keratocytes–specialized fibroblasts in the cornea that elongate their cell bodies between lamellae of collagen fibrils–and the corneal mesenchymal stromal cells (MSCs) from which the keratocytes are derived [59,60]. Corneal MSCs can be readily harvested and expanded, and they are known to play a crucial role in regenerating corneal tissue after damage or disease [61–64]. Therefore, collagen gels laden with corneal MSCs may have potential to address the global shortage of donor corneas required for allograft transplantation to treat corneal blindness. These corneal MSCs remain highly viable (>90% viability) over 5 days in culture encapsulated within PHYS, SPAAC, and IPN collagen gels (**Figure S7**).

The forces that contractile cells exert on their surrounding matrix can induce significant changes in the overall shape and size of the hydrogel. In particular, a challenge of conventional collagen gels is the severe densification and contraction in size that occurs as contractile cells pull on physically self-assembled collagen fibrils [7–9]. We observed that the different crosslinking mechanisms and network microstructures of PHYS, SPAAC, and IPN collagen affected the extent to which these hydrogels contracted (Figure 5). All gels were prepared in circular 4-mm diameter molds with an initial concentration of 0.5 million corneal MSCs/mL, and the sizes of the gels were tracked daily over 5 days of culture. Due to the brightfield imaging of the gels, the fibrillar PHYS and IPN collagen gels appear darker than the amorphous SPAAC collagen gels on Day 0, since light scattering from fibrils causes opacity [65]. Over the 5-day culture period, the PHYS collagen gels with encapsulated corneal MSCs were rapidly and severely contracted. On average, PHYS collagen gels contracted to <30% of their initial gel area within 2 days and <10% of their initial gel area within 4 days of cell encapsulation. Furthermore, we observed that the PHYS collagen gels became increasingly opaque over time, consistent with collagen fiber densification during contraction [65]. In contrast, the material conditions with amorphous SPAAC collagen in the network—both SPAAC collagen alone and the IPN collagen—did not detectably contract in response to the encapsulated corneal MSCs, indicating that the amorphous, covalently crosslinked collagen network significantly increases the stability of the collagen hydrogels against cell-induced contraction.

**Figure 5.**
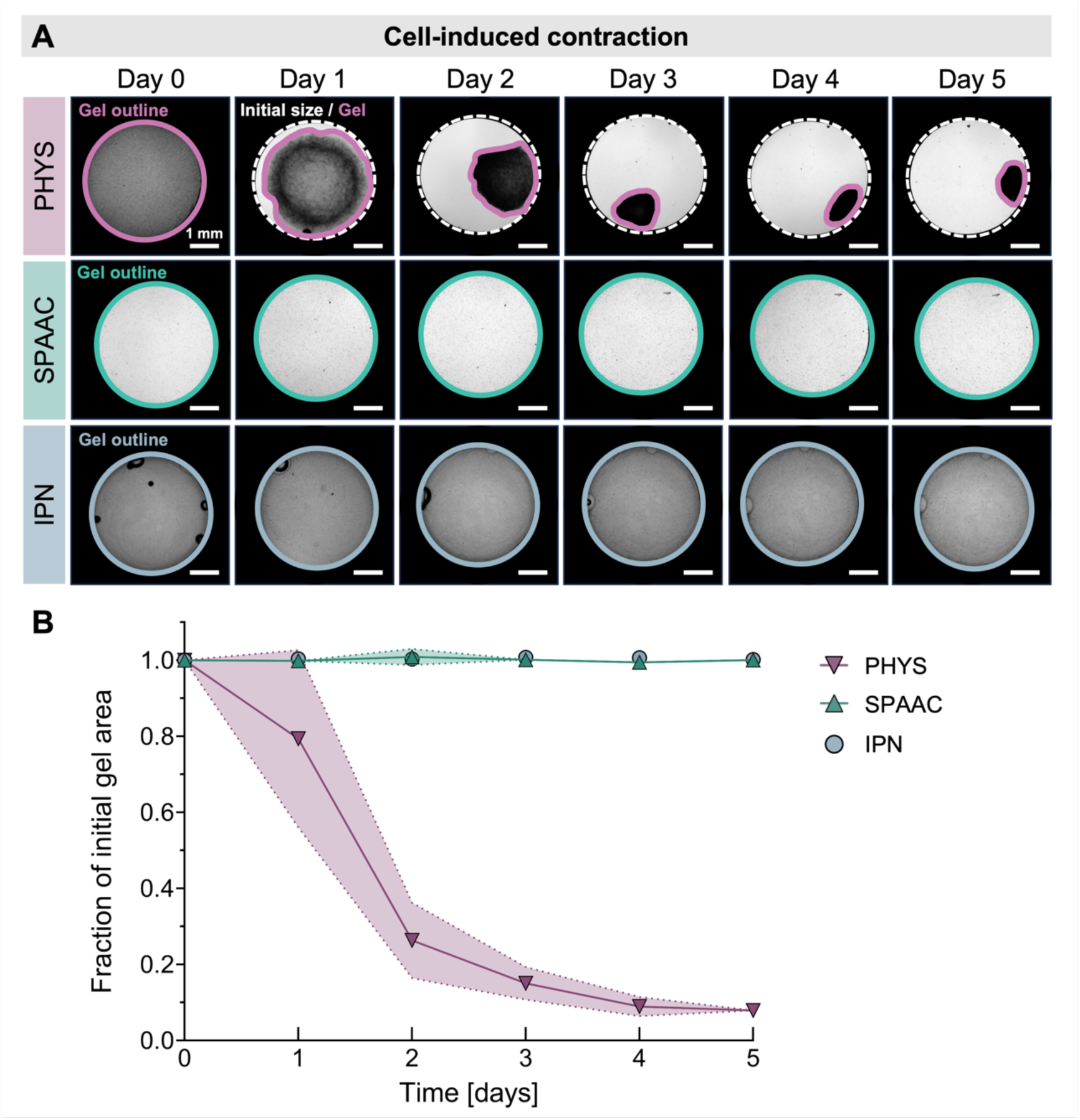
Stability of cell-laden PHYS, SPAAC, and IPN collagen gels against cell-induced contraction. **(A)** Representative brightfield images of gel shapes over time compared to the initial gel size. PHYS collagen with 0.5 million corneal MSCs/mL experienced severe contraction over 5 days in culture, while SPAAC and IPN collagen with 0.5 million corneal MSCs/mL did not detectably contract from their initial gel size. **(B)** Rate of collagen gel contraction. N ≥ 5 independent gels per material condition. Shaded regions represent the standard deviation from the mean.

### 3.4. Morphology of cells encapsulated in the collagen IPN

The morphology of mammalian cells is a key indicator of their function [66,67]. Therefore, understanding how cell spreading is regulated by microenvironmental cues is important for the design of biomaterials for tissue engineering [53,68,69]. In native corneal tissue, the corneal MSCs spread and elongate within the collagen-rich stroma [61,63]. We hypothesized that the presence of collagen fibrils within our IPN collagen may allow for cell spreading (Figure 6), even within a stable gel that does not experience bulk contraction (Figure 5). Indeed, corneal MSCs were able to spread in both PHYS and IPN collagen hydrogels that contained fibrils, elongating their cell bodies. In contrast, cells in the amorphous, covalently crosslinked SPAAC collagen alone remained rounded, extending only small protrusions (Figure 6A). While the extent of cell spreading in the collagen IPN was initially slightly less than in PHYS collagen alone (on Days 0 and 1), the cells were able to reach and maintain the same degree of spreading in the collagen IPN as in PHYS collagen within a short timespan, by Day 3 in culture (Figure 6B).

**Figure 6.**
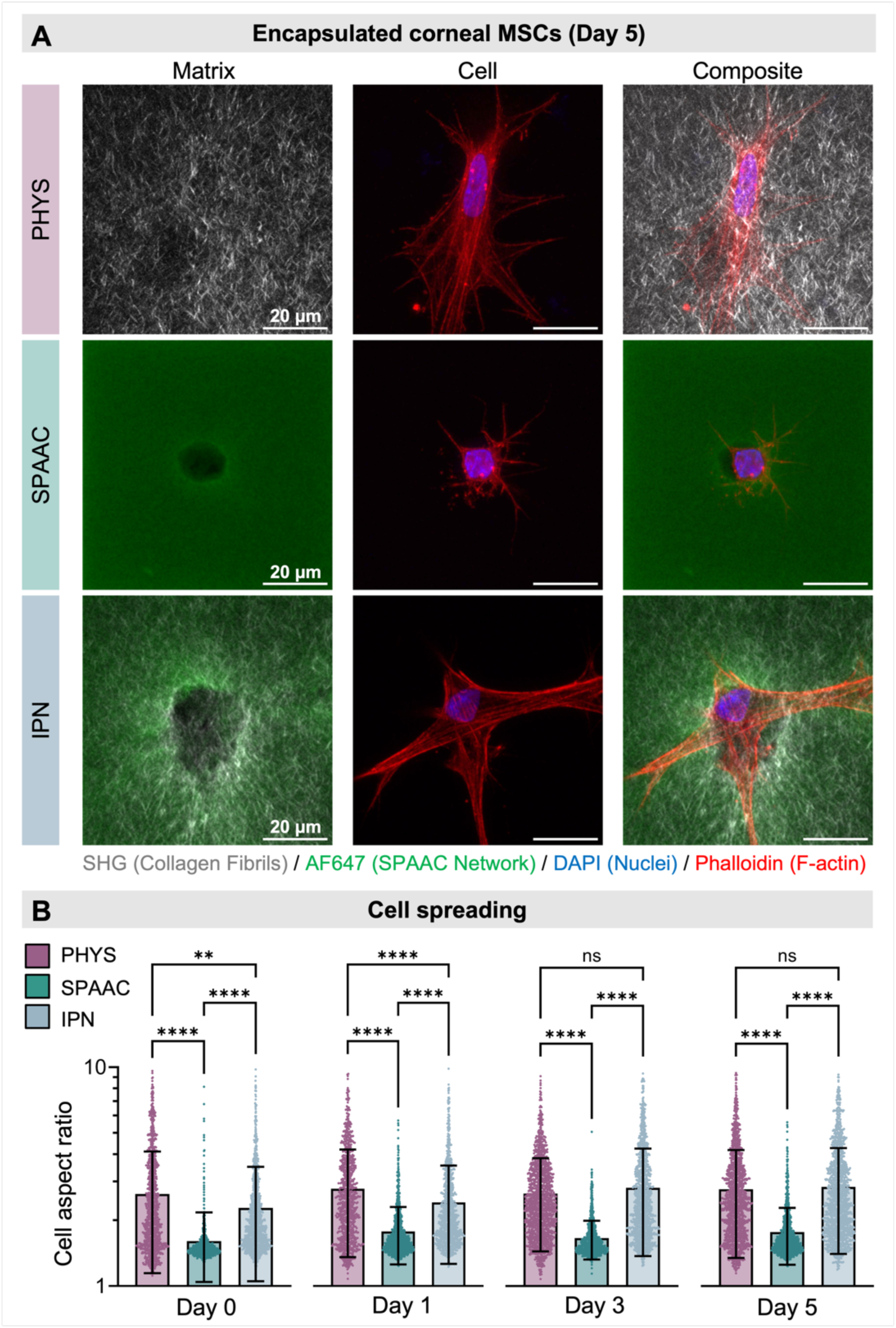
Cell spreading behavior and matrix morphology. **(A)** Representative composite SHG and fluorescence microscopy images show extensive spreading of corneal MSCs in the PHYS and IPN collagen gels by Day 5 after cell encapsulation, with significantly reduced cell spreading in the SPAAC gels. **(B)** Quantification of the aspect ratios of cells in PHYS, SPAAC, and IPN collagen gels throughout the cell culture period. Statistical analyses were performed with the Kruskal-Wallis nonparametric test and Dunn’s multiple comparisons correction. In all cases, N = 3 independent gels and n = 500 - 4000 cells per material condition and time point. Data plotted as mean ± SD. ns = not significant, ** p < 0.01, **** p < 0.0001.

By imaging the fibrillar collagen IPN around the corneal MSCs over time, we observed local densification and realignment of collagen fibrils near cellular projections. The local reorganization of the collagen appeared within 24 h but then remained seemingly stable over time (**Figure S8**). Corneal MSCs can therefore interact with and reorganize the collagen fibril networks in IPN collagen as the cells spread. However, while the local densification of the collagen fibrils around the cells for PHYS and IPN collagen was similar, the long-range densification of collagen that results in severe bulk contraction of the gel was observed only for PHYS collagen and not for IPN collagen (Figure 5). In fibrillar matrices in which the fibrils are less connected to each other, forces propagate in fewer directions and with less robust force transmission compared to highly connected networks [70]. Despite PHYS and IPN collagen having similar fibrillar microstructures (Figure 3), we expect that the covalently crosslinked, SPAAC collagen network that fully surrounds the fibrils in the collagen IPN likely serves as a barrier between fibrils that dampens transmission of forces and inhibits long-range fibril rearrangement, thus preventing the gel from contracting. In this way, the collagen IPN allows cells to interact with collagen fibrils on a microscopic scale and adopt spread morphologies, while also maintaining a highly stable and non-contracting bulk gel structure on a macroscopic scale.

## 4. Discussion

A key challenge in tissue engineering is the tradeoff between allowing cells to remodel their ECM and spread while preserving the overall structural stability of the gels. Maintaining the shape and size of cell-laden hydrogels is critical for applications in personalized regenerative medicine, where precise dimensions and structural stability are necessary for implant success. This is especially challenging for tissue engineered constructs with contractile cells such as MSCs, which can exert significant forces on their surrounding matrix as they spread, leading to severe contraction of the gel. Herein, we described our development of a collagen-based IPN that offers the benefits of both physical self-assembly of collagen (to form a fibrillar microstructure that the cells can interact with and realign on a microscopic scale) as well as covalent crosslinking of collagen (to stabilize the bulk hydrogel on a macroscopic scale). Both networks within our IPN are based on collagen, but the different crosslinking approaches for the two networks (PHYS and SPAAC collagen) allow for a unique microstructure consisting of collagen fibrils embedded within an amorphous, covalently crosslinked collagen matrix, thus promoting both cell spreading and structural integrity simultaneously.

Our results indicate that the spreading of the human corneal MSCs is facilitated by the presence of a fibrillar collagen network. Interestingly, the cells were able to spread in the collagen IPN despite the simultaneous presence of the amorphous SPAAC collagen network, in which the corneal MSCs do not spread. It is possible that the chemical crosslinking of SPAAC collagen may also obscure some cell binding sites [71]. To better understand the ability of the collagen IPN to promote cell spreading while maintaining structural integrity, our findings motivate future investigations of the propagation of a force exerted on a collagen fiber within an amorphous, covalently crosslinked network. In particular, our current results could be supplemented with computational modeling that considers cell-generated forces, substrate deformation, ECM density, cell migration, and proliferation [72–74]. Unlike most current models of cell-generated forces on collagen fibers, our collagen IPN is not solely fibrillar, but rather has amorphous, covalently crosslinked SPAAC collagen filling the voids between fibrils. The ability of corneal MSCs to locally rearrange collagen fibrils and spread within the collagen IPN indicates that short-range interactions between cells and collagen fibrils remain intact, facilitating essential cellular processes. Since this does not induce widespread collagen IPN gel contraction, long-range force propagation is likely inhibited by the amorphous, covalently crosslinked SPAAC collagen gel occupying the interfibrillar space, which stabilizes the overall gel structure.

Looking forward, this collagen IPN could be adapted to a variety of applications for both tissue model systems *in vitro* as well as implantable therapies *in vivo*. Future development could vary the concentrations and ratios of PHYS and SPAAC collagen to tune material properties of the IPN based on the cell type and the downstream application of the biomaterial. Furthermore, we expect that the collagen IPN could be adapted as an ink for embedded 3D bioprinting, with diffusion of the PEG-DBCO crosslinker from the support bath into the print [11,75,76]. This would expand possibilities for creating patient-specific collagen implants with tunable properties. Overall, this collagen IPN offers a versatile foundation for developing hydrogel scaffolds that bridge the gap between cellular functionality and material stability to improve the efficacy and reliability of bioengineered collagen hydrogels.

## 5. Conclusions

In summary, we have developed an IPN based on two collagen networks that effectively integrates physical self-assembly (fibrillar, PHYS collagen) with covalent crosslinking (amorphous, SPAAC collagen). The unique crosslinking combination and microstructure of the collagen IPN promotes spreading of encapsulated cells without compromising bulk gel stability. By sequentially forming the IPN with (1) fibrillar, PHYS collagen and then (2) amorphous, SPAAC collagen to fill the interfibrillar space, we achieved a fibrillar microstructure similar to PHYS collagen alone. Importantly, this IPN was able to facilitate the spreading of encapsulated human corneal MSCs (like PHYS collagen alone) with negligible amounts of bulk gel contraction (like SPAAC collagen alone). Overall, our results suggest that this engineered collagen IPN holds significant potential for tissue engineering applications by promoting key cellular processes like spreading while preserving scaffold shape and stability.

## Data availability

All data needed to evaluate the conclusions in the paper are present in the paper and/or the Supplementary Information.

## Acknowledgements

The authors thank Dr. Ninna Rossen for helpful discussions about the Fiber Finding Analysis. The authors acknowledge funding support from the National Science Foundation including DGE-165618 (L.G.B., C.M.L.), DMR-2103812 (S.C.H.), DMR-2427971 (S.C.H.), and CBET-2033302 (S.C.H); the National Institutes of Health including F31-EY034785 (L.G.B), P30-EY026877 (D.M.), R01-EY033363 (D.M.), and R01-EY035697 (D.M., S.C.H.); the ARCS Foundation Scholarship (L.G.B.); the Stanford Bio-X Interdisciplinary Graduate Fellowship (C.M.L., B.C.); the Swiss National Science Foundation including P500PN210723 (F.C.); the Stanford Knight-Hennessy Scholars Program (B.C.); the Stanford Graduate Fellowship in Science and Engineering (D.S.); a departmental core grant from Research to Prevent Blindness (D.M.), and the Advanced Research Projects Agency for Health including ARPA-H HEART (S.C.H).

