## Supplementary Information for "Interpenetrating networks of fibrillar and amorphous collagen promote cell spreading and hydrogel stability"

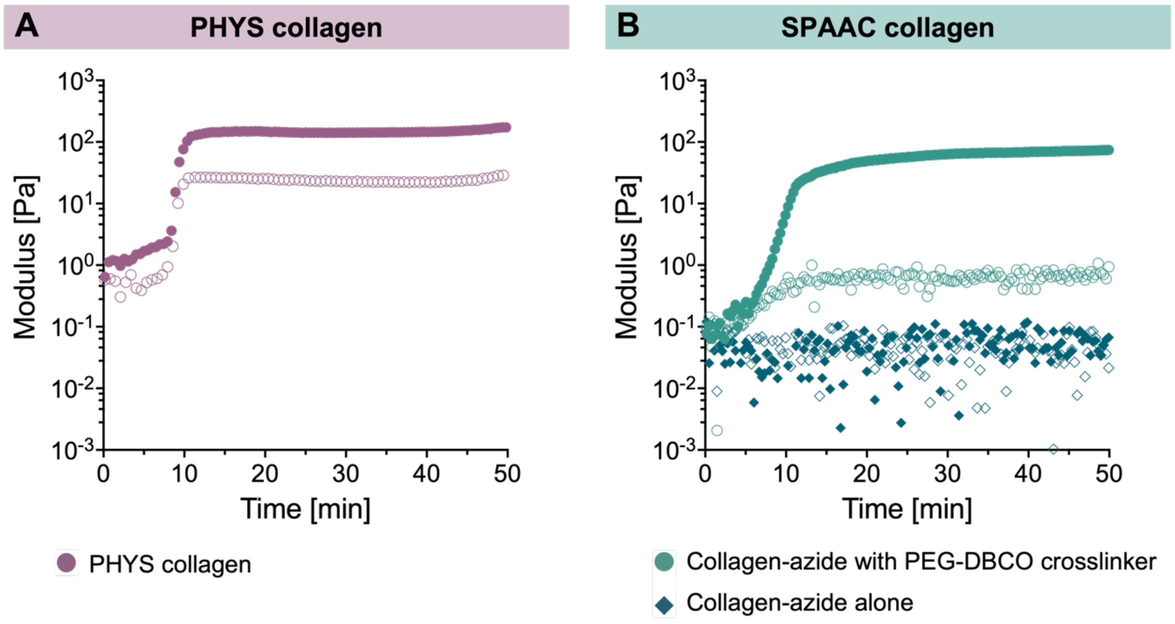


**Figure S1.** Gelation properties of PHYS and SPAAC collagen networks. Filled symbols represent the storage modulus (G'), and open symbols represent the loss modulus (G"). **(A)** PHYS collagen gelates spontaneously at a neutral pH (~7.5) and 37°C within 10 min. **(B)** Collagen-azide does not spontaneously gelate at 37°C on its own. When collagen-azide is mixed with the strained alkyne PEG-DBCO crosslinker, the SPAAC reaction proceeds to covalently crosslink the gel within 10 min.

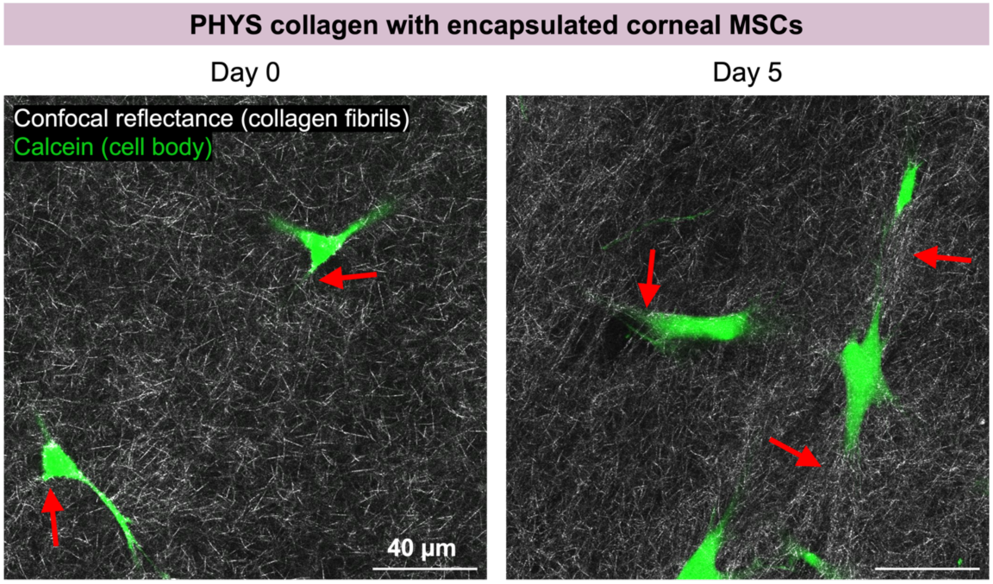


**Figure S2.** Representative confocal reflectance images of PHYS collagen gels with encapsulated corneal MSCs. The collagen fibrils appear more compacted and aligned in the pericellular region near cellular projections (red arrows). This local ordering of collagen fibrils is more pronounced at Day 5 versus Day 0.

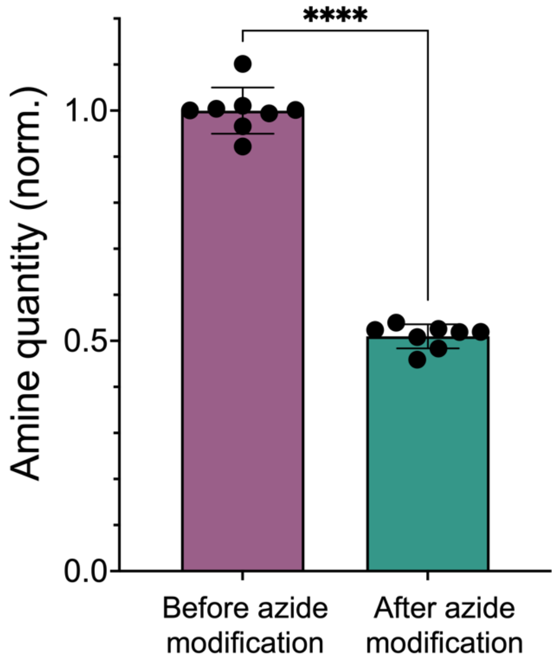


**Figure S3.** During the collagen-azide bioconjugation reaction, approximately half of the primary amines in the collagen are modified with azides through N-hydroxysuccinimide (NHS) ester chemistry. Normality of the data was confirmed with the Shapiro-Wilk test, and statistical analysis was performed with an unpaired t test. N = 8 independent samples per material condition. Data plotted as mean ± SD. **** $p < 0.0001$.

**Table S1.** Statistical comparison of fibril network properties between PHYS collagen, the IPN collagen with simultaneous gelation of the PHYS and SPAAC networks, and the IPN collagen with sequential gelation of the PHYS and SPAAC networks. N = 3 independent gels per material condition. Statistical analyses performed as one-way analysis of variance (ANOVA) with Tukey's multiple comparisons test.

| | **PHYS vs. IPN, simultaneous** | **PHYS vs. IPN, sequential** | **IPN, simultaneous vs. IPN, sequential** |
|---|---|---|---|
| **Contour length** | <0.0001 (****) | <0.0001 (****) | 0.0016 (**) |
| **Persistence length** | <0.0001 (****) | 0.0599 (ns) | <0.0001 (****) |
| **Mesh size** | <0.0001 (****) | <0.0001 (****) | <0.0001 (****) |
| **Fiber count** | <0.0001 (****) | <0.0001 (****) | <0.0001 (****) |
| **Fiber width** | <0.0001 (****) | 0.0052 (**) | <0.0001 (****) |

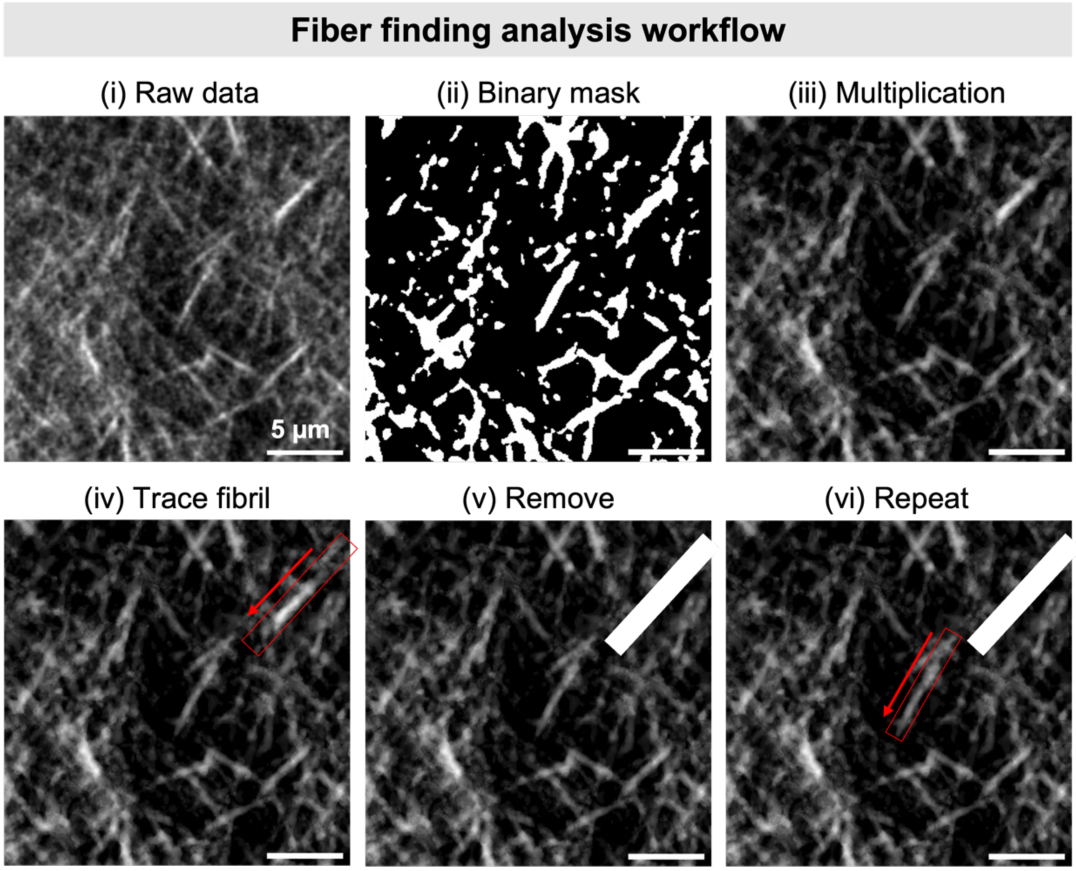


**Figure S4.** Fiber finding analysis workflow for second harmonic generation (SHG) images. Using FIJI, raw signal data is thresholded into a binary mask that is multiplied into the original image to remove background noise. Subsequently, using a MATLAB algorithm developed by Rossen et al., each fibril is traced and blotted out from the image in an iterative fashion until all fibrils have been quantified.

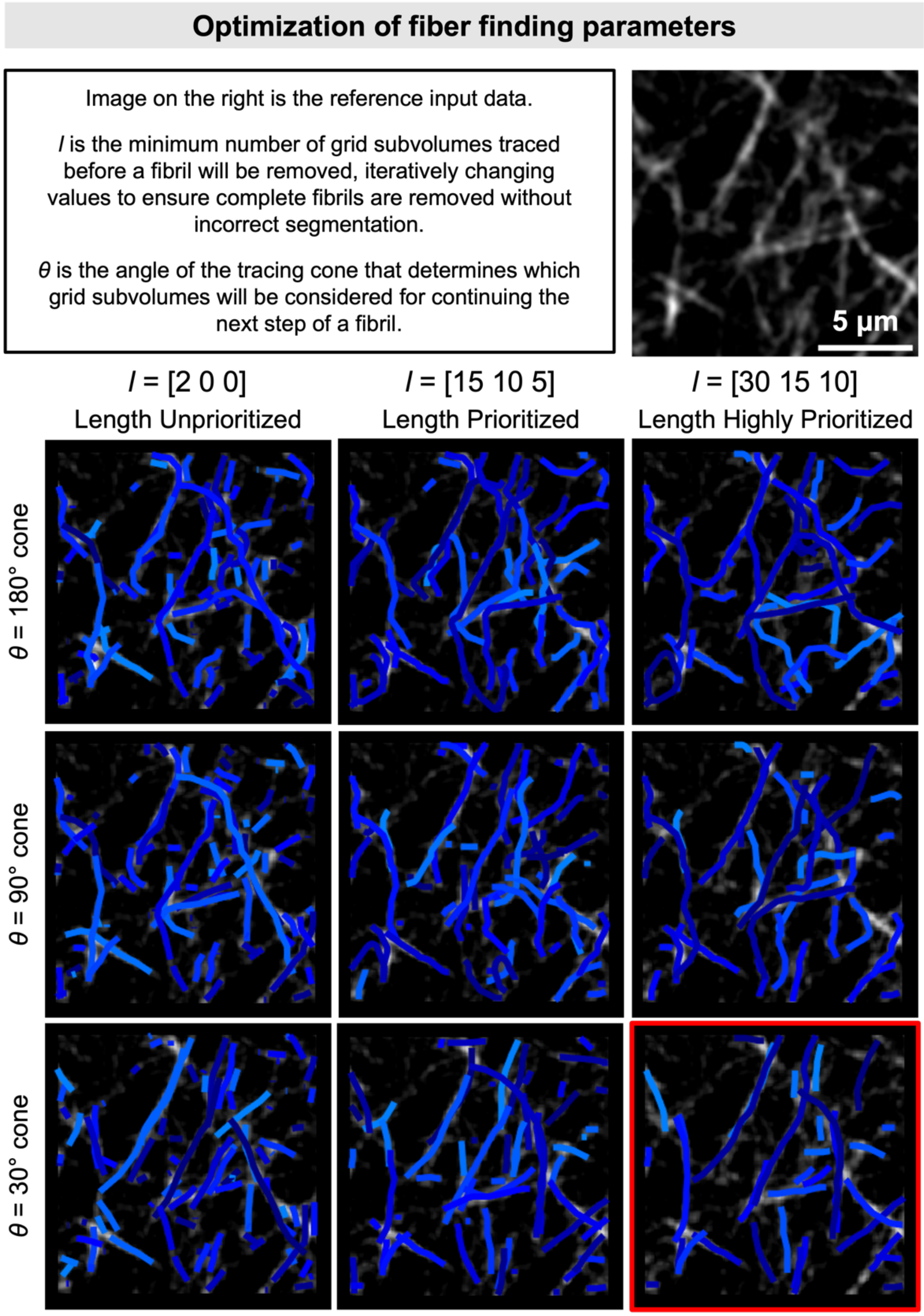


**Figure S5.** Parameter optimization scheme for fiber finding analysis algorithm. Tracing cone angle and fibril length prioritization array were parametrically optimized to select a set of variables that yields the most accurate fibril identification based on visual inspection.

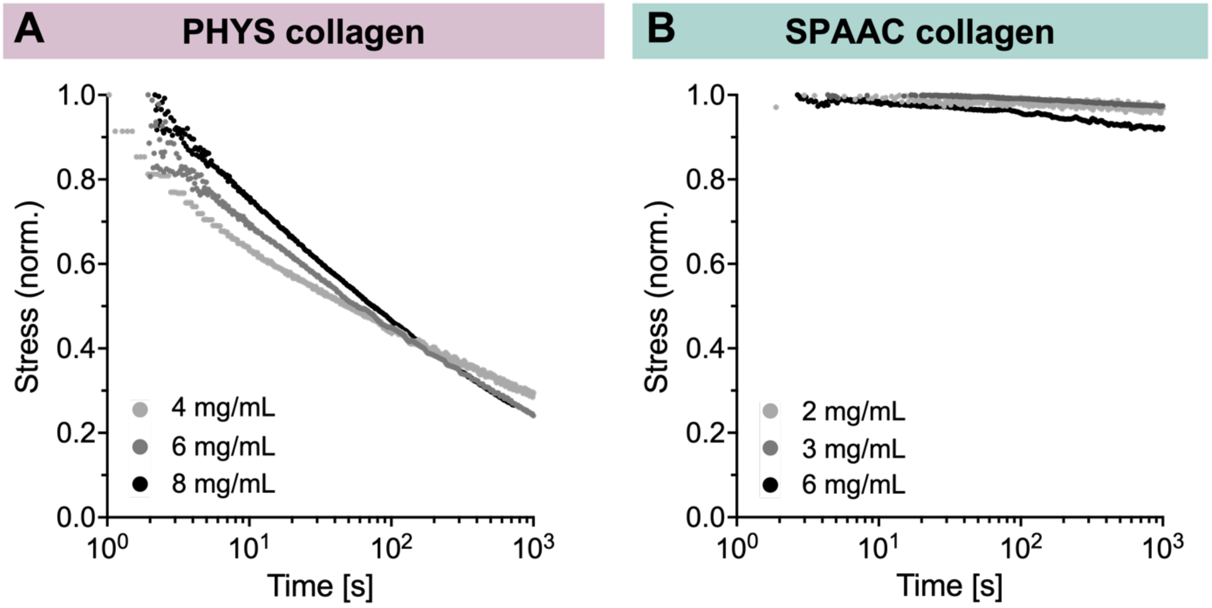


**Figure S6.** Representative stress relaxation tests on gels made from **(A)** PHYS collagen and **(B)** SPAAC collagen indicate consistent stress-relaxation behavior for each type of network regardless of collagen concentration.

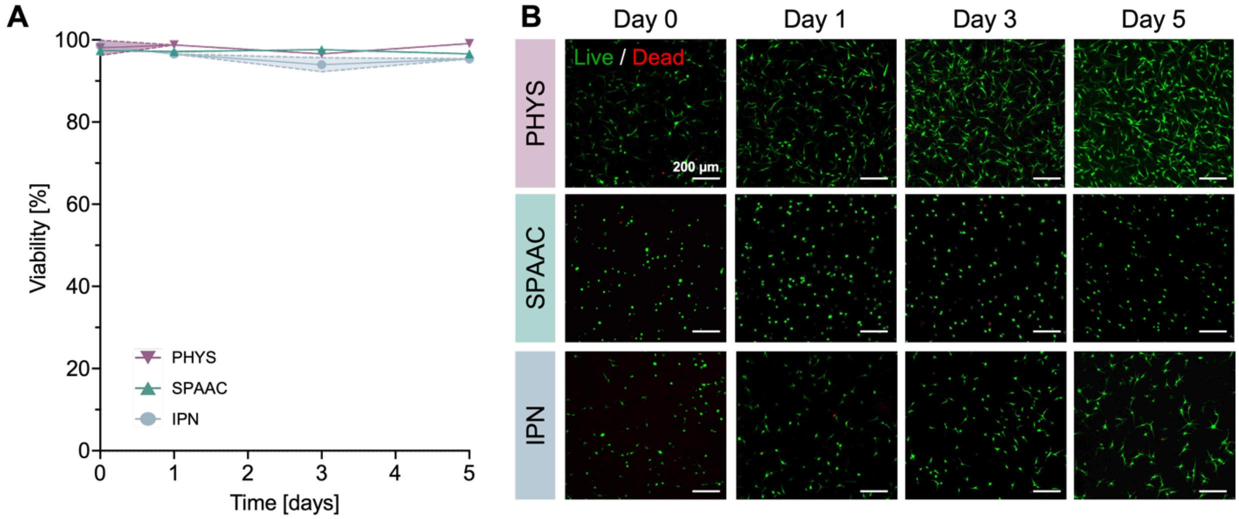


**Figure S7.** Corneal MSC viability in 3D collagen gels. **(A)** Corneal MSCs encapsulated in PHYS, SPAAC, and IPN collagen remain highly viable over 5 days in culture. **(B)** Representative images from Live / Dead cytotoxicity assays of encapsulated corneal MSCs on the day of encapsulation (Day 0) and on Days 1, 3, and 5 after encapsulation. N = 3 independent gels per material condition and time point. Shaded regions represent the standard deviation from the mean.

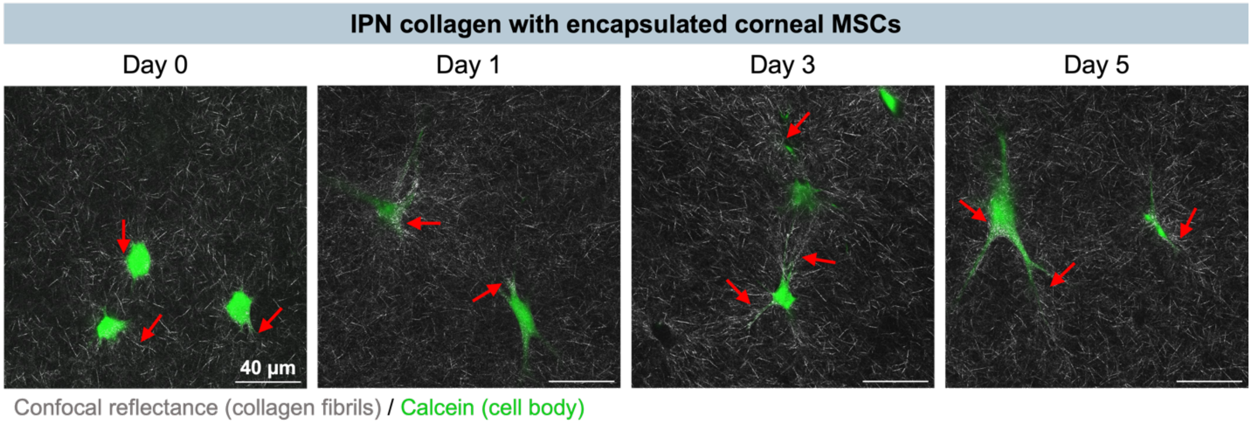


**Figure S8.** Representative confocal reflectance and fluorescence images showing corneal MSC interactions with the collagen matrix in the IPN collagen hydrogels. Local densification and alignment of collagen fibrils to the cellular projections occurs within 24 hours and is seemingly stable over time.
